# An Integrative Phylogenomic Framework Quantifies the Dominant Role of Introgression in Phylogenetic Discordance among Diploid *Oryza* Species

**DOI:** 10.64898/2026.02.01.702050

**Authors:** Hui-fang Li, Shuang-feng Dai, Tian-long Fang, Li-zhi Gao

## Abstract

**Background:** Phylogenomic studies frequently reveal widespread gene tree discordance, primarily arising from incomplete lineage sorting (ILS) and hybridization and/or introgression. Disentangling these processes is especially challenging in rapidly radiating lineages. The genus *Oryza*, with its rapid diversification and multiple genome types, exemplifies this pervasive phylogenetic incongruence. We integrated multiple genomic datasets including whole-genome resequencing, transcriptomes, and published genomes from diploid *Oryza* species. Concatenation and multispecies coalescent analyses recovered a robust, congruent species tree, placing the FF and GG genome groups as a monophyletic basal clade, followed by successive divergence of the EE, CC, BB, and AA lineages, a topology differing from some prior hypotheses.

**Results:** To assess the sources of discordance, we employed a suite of complementary phylogenomic methods. Quantifying introgression via Branch Lengths (QuIBL)-based model comparisons suggested that ∼74.17% of gene tree-species tree discordance is better explained by post-speciation introgression, whereas only ∼15.56% is consistent with ILS alone. Phylogenetic networks (PhyloNet) and allele-sharing statistics (*D*-statistics, *f*-branch) corroborated these results, indicating widespread historical introgression both within and between genome groups. Furthermore, genome-wide scans using the *f*_dM_ statistic localized introgressed genomic regions, which showed reduced interspecific divergence and were enriched for genes involved in stress responses and metabolism.

**Conclusions:** Taken together, our results demonstrate that historical introgression, not ILS, is the dominant force shaping phylogenetic discordance in diploid *Oryza*. The integrative phylogenomic framework implemented here, which quantifies the contributions of introgression versus ILS and maps the genomic footprint of gene flow, provides a replicable strategy for resolving complex evolutionary histories in other rapidly radiating lineages.

## Background

The evolutionary history of many biological lineages is shaped not by strictly bifurcating trees but by reticulate processes, such as hybridization and/or introgression, which facilitate genetic exchange across species boundaries [1, 2]. These processes frequently lead to discordance between gene trees and the species tree, a pervasive phenomenon known as phylogenetic incongruence [3]. ILS, introgression, and adaptive radiation are all primary drivers of such discordance [4–7]. In addition, pronounced heterogeneity in genome-scale evolutionary processes can result in distinct evolutionary histories among different genes or genomic regions [8]. This complexity is compounded during rapid radiations, where ILS can obscure signals of introgression, making it particularly challenging to reconstruct an accurate evolutionary history [9–13].

The rise of phylogenomics has greatly advanced the reconstruction of the Tree of Life, with genome-scale data and novel statistical methods resolving long-standing controversies in various lineages [14–19]. However, the increasing use of multilocus data has also revealed that phylogenetic conflict, especially gene tree-species tree discordance, is widespread. In plants, incongruence has been documented at multiple taxonomic levels, from within genera [20–25], among genera within families [26–29], and even among orders or major angiosperm lineages [30–33]. Accurately inferring evolutionary relationships requires distinguishing whether such conflict stems from genuine biological processes, like ILS or introgression [34, 35], or from analytical artifacts [36], a challenge that persists despite methodological advances [37–41].

Significant progress has been made in developing tools to address discordance. Species tree methods like ASTRAL [42], SNAPP [43], and SVDquartets [44] explicitly account for ILS, while network-based approaches like PhyloNetworks [45] model reticulate evolution. Nevertheless, even genome-scale phylogenies with high support can harbor extensive underlying gene tree conflict. To dissect this conflict, quantitative methods such as the *D*-statistic [46, 47] and QuIBL [48] have been developed to detect introgression and distinguish it from ILS, successfully illuminating complex histories in groups like cichlids and *Heliconius* butterflies [45, 48]. Disentangling these biological factors from error is therefore crucial for understanding speciation and adaptation, especially in groups where hybridization may also trigger polyploidization.

The genus *Oryza* (Poaceae), encompassing cultivated rice and wild relatives, is a system of great agricultural and evolutionary importance, and an ideal model for studying the causes of gene tree conflict. It includes 27 species with 11 genome types: six diploid (AA, BB, CC, EE, FF, GG) and five allopolyploid (BBCC, CCDD, HHJJ, HHKK, KKLL) types [49–51]. The genus has a clear history of reticulation; all tetraploids are allopolyploids derived from diploid hybrids [52–55], including BBCC types formed from BB and CC genome parents and the CCDD genome involving a CC maternal and EE (*O. australiensis*) paternal parent [56].

Phylogenetic studies in *Oryza* have long been marked by incongruence. Early work using few loci consistently reported strong gene tree conflict regarding relationships among major genome types [23, 56–63]. For instance, Ge et al. [23] found chloroplast (*matK*) and nuclear (*Adh1*, *Adh2*) trees supported conflicting relationships among AA, BB, and CC genomes, and disagreed on whether FF and GG genomes diverged sequentially (nuclear) or formed a clade (chloroplast). While relationships within the AA genome are now relatively clear with *O. rufipogon* and *O. barthii* identified as progenitors to Asian and African cultivated rice, respectively [50, 63], and *O. meridionalis* and *O. longistaminata* occupying basal positions [50, 62, 63], the deep branching order, particularly the positions of the FF and GG genomes, remains contentious. Some studies suggest GG is the most basal diploid lineage, followed by FF [51, 59], while chloroplast data often group FF and GG together [61, 63]. This persistent discordance, evident even in genome-scale analyses [50, 64–67], highlights the complex interplay of ILS and introgression in the genus’s history and the challenge of using multilocus phylogenetic analyses to resolve such conflicts [68]. However, previous research has primarily documented the presence of gene flow; a comprehensive, quantitative assessment partitioning the relative contributions of introgression versus ILS to genome-wide phylogenetic discordance in *Oryza* is still lacking.

In this study, we leverage multiple genomic datasets including whole-genome resequencing, transcriptomes, and published genomes to achieve three main objectives. First, we aim to reconstruct a robust species phylogeny for diploid *Oryza* using both concatenation and multispecies coalescent methods applied to multi-omics data. Second, and central to our approach, we develop and apply an integrative phylogenomic framework designed to quantify the relative contributions of ILS and introgression to genome-wide gene tree conflict. This framework employs complementary methods including phylogenetic networks (PhyloNet), allele-frequency statistics (*D*-statistic, *f*-branch), and model-based comparison of gene tree branch lengths (QuIBL) to distinguish between these two primary sources of incongruence. Third, we localize introgressed genomic regions and assess their potential functional significance. By integrating these analyses, we aim to resolve long-standing phylogenetic uncertainties in *Oryza* and to elucidate the predominant evolutionary forces that have shaped the history of this critical plant genus.

## Results

### Sequencing data overview and quality assessment

Our dataset comprised whole-genome resequencing data for two individuals from each of the 16 diploid *Oryza* genome types, plus the outgroup *Leersia perrieri* (33 accessions total, Additional file 1: Table S1). All samples were aligned to the *O. sativa* cv. Nipponbare reference genome to identify high-quality SNPs. Principal component analysis (PCA) based on genome-wide SNPs confirmed that individuals from the same species clustered together, and species grouped by genome type, indicating high data reliability and biological consistency (Additional file 2: Fig. S1a). Except for *O. rufipogon*, the two individuals per species yielded comparable SNP counts (Additional file 2: Fig. S1b); the larger discrepancy in *O. rufipogon* is likely due to its high intraspecific genetic diversity, though both samples still clustered closely in PCA.

SNP counts generally increased with phylogenetic distance from the reference. However, EE, FF, and GG genome species had fewer SNPs than cultivated rice, a pattern likely caused by reduced mapping efficiency due to greater sequence divergence (Additional file 2: Fig. S1b). SNP distribution varied markedly among species at the chromosomal level, reflecting extensive genetic diversity. For example, within a specific interval on chromosome 11 (25,700,000-25,800,000 bp), CC genome species harbored substantially more SNPs than others (Additional file 2: Fig. S1c).

After stringent filtering, 187,135 SNPs at 4DTv sites and 209,888 SNPs within single-copy gene regions were retained for phylogenetic analysis (Additional file 2: Fig. S1d, Additional file 1: Table S2). These SNPs were distributed across all 12 chromosomes, predominantly in upstream, downstream, and exonic regions.

To complement genomic data and validate orthology, particularly for species lacking high-quality reference genomes, we generated *de novo* transcriptome assemblies for all 16 diploid species (Additional file 1: Table S3). BUSCO assessments indicated high completeness (∼90%) (Additional file 2: Fig. S2), and we identified 846 single-copy orthologous genes from these assemblies (Additional file 1: Table S3). Furthermore, protein sequences from 13 publicly available diploid genomes yielded 3,973 single-copy orthologs, providing an independent dataset for phylogenetic validation and collectively confirming the robustness of our evolutionary inferences.

### A robust phylogenetic framework for diploid *Oryza*

We applied concatenation-based and coalescent-based phylogenomic approaches to nuclear and plastid datasets to establish a reliable species tree. Phylogenies based on SNPs from single-copy gene regions and 4DTv sites were highly congruent (Fig. 1a, b) and consistently resolved two major clades: a basal clade containing the FF and GG genome types, and a second clade comprising the EE, CC, BB, and AA genome groups (Fig. 1a, b). Within the AA genome group, *O. meridionalis* was basal, followed by *O. longistaminata* and *O. glumaepatula*. *O. glaberrima* clustered with its wild progenitor *O. barthii*, and cultivated indica rice (*O. sativa*) grouped with *O. nivara* and *O. rufipogon* (Fig. 1a, b). As *indica* accessions were used, *O. sativa* clustered more closely with *O. nivara*, consistent with previous findings that *O. nivara* represents the direct ancestor of *indica* rice, whereas *O. rufipogon* is the progenitor of *japonica* rice [50].

**Fig. 1.**
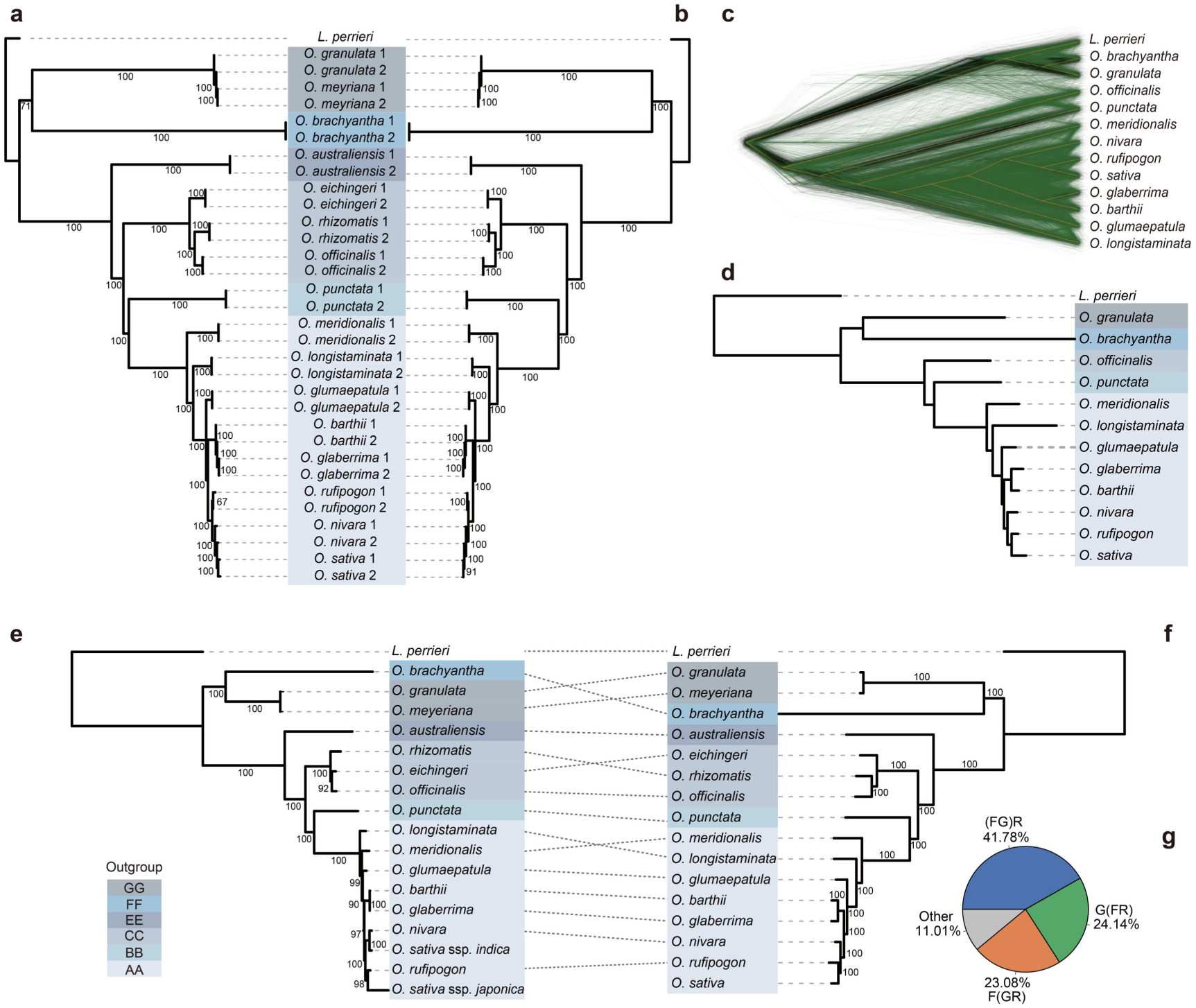
Phylogenetic reconstruction of diploid *Oryza* species. **a** ML tree inferred from SNPs within single-copy orthologous gene regions. **b** ML tree inferred from SNPs at 4DTv sites. **c** Visualization of nuclear gene tree discordance using DensiTree. The consensus species topology (yellow line), which is congruent with the tree in panel d, is shown against the cloud of individual gene tree topologies (green and black lines). **d** Concatenated ML phylogeny reconstructed from 3,973 single-copy orthologous genes derived from genomic data. **e** Chloroplast genome-based ML phylogeny. **f** Concatenated ML phylogeny reconstructed from 846 single-copy orthologous genes derived from transcriptome assemblies. **g** Frequency distribution of topologies among individual gene trees from the transcriptomic dataset. “R” represents topologies where lineages other than FF or GG occupy the basal position.

This topology, with FF and GG as a joint basal clade, differs from some previous studies that suggested sequential divergence of GG then FF [23, 51, 59]. Our finding was strongly corroborated by independent datasets: a concatenated tree from 3,973 genomic orthologs (Fig. 1d), a coalescent-based species tree from the same genes (Fig. 1c), and a phylogeny from 846 transcriptome-derived orthologs all recovered the same relationship (Fig. 1f). Analysis of individual gene trees showed 41.78% supported the FF+GG basal clade, a proportion significantly higher than those supporting FF (23.08%) or GG (24.14%) alone (Fig. 1g). The concordance between datasets underscores the robustness of the inferred relationships and aligns with observations that individual gene trees do not necessarily reflect the species tree [59], yet concatenation of multiple loci can maximize phylogenetic information.

To further test the nuclear framework, we reconstructed a plastid phylogeny. Given that plastid genomes evolve slowly and exhibit conserved structure, making them widely used in deep-level plant phylogenetics [69–71], the plastid tree also placed FF and GG as a sister clade to all other diploid genomes (Fig. 1e). However, cytonuclear discordance was observed within the AA lineage: the plastid tree placed *O. longistaminata*, not *O. meridionalis*, as the earliest-diverging lineage (Fig. 1e), suggesting historical chloroplast capture.

In summary, analyses across multiple datasets consistently demonstrate that FF and GG genomes form a distinct basal clade. The widespread topological discordance indicates that extensive hybridization and/or ILS have shaped diploid *Oryza* evolution.

### Genome-wide patterns of gene tree discordance

Despite a consistent species tree, we detected extensive gene tree conflict. PhyParts analysis of 3,973 nuclear gene trees mapped onto the species tree revealed widespread discordance (Fig. 2a). While 1,741 genes (43.82%) supported the FF+GG basal clade, conflict was high at shallower nodes. For example, only 19.03% of gene trees agreed with the species tree topology for relationships among Asian (*O. rufipogon*, *O. nivara*, *O. sativa*) and African (*O. barthii*, *O. glaberrima*) AA genome species.

**Fig. 2.**
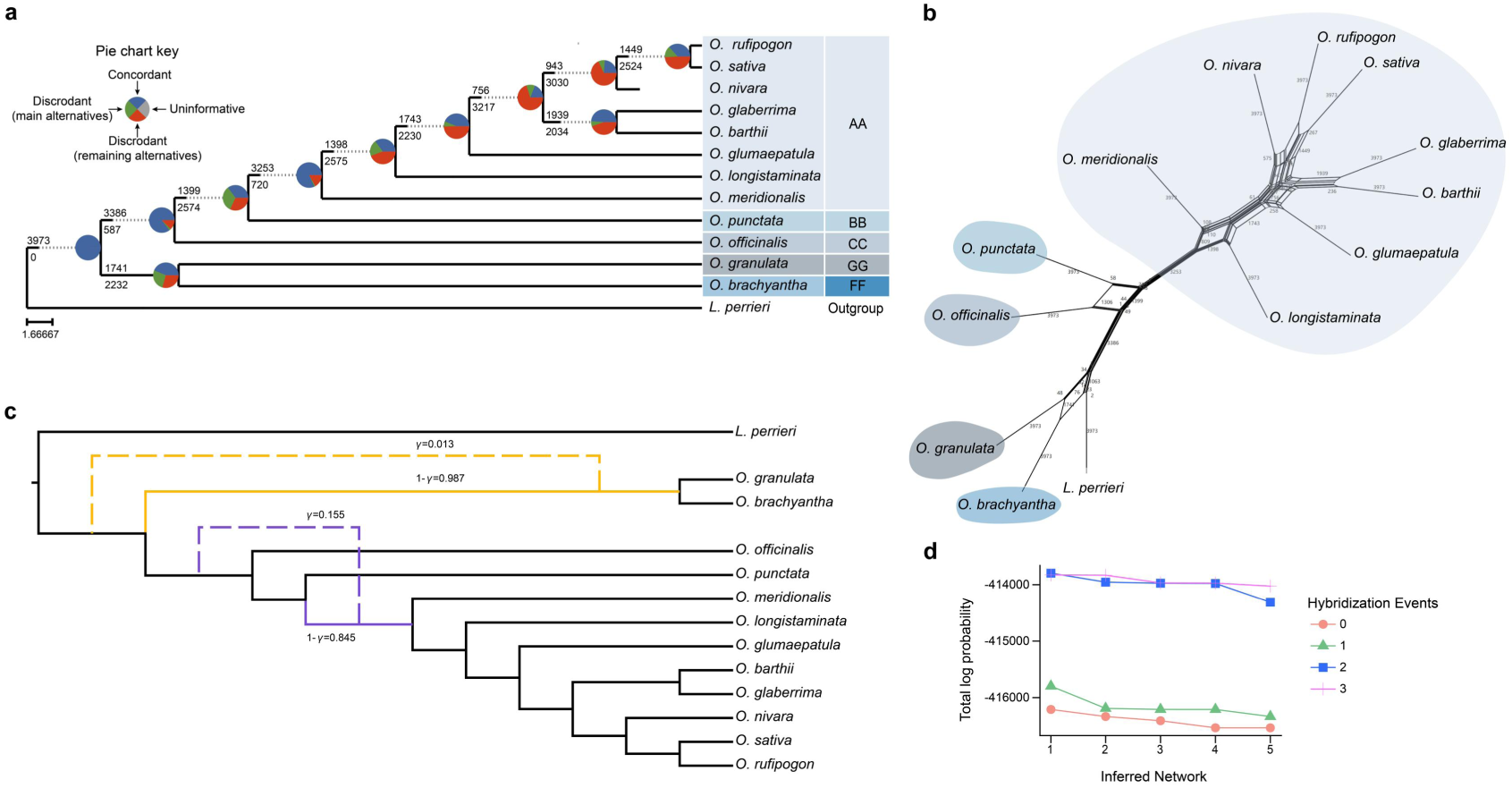
Concordance, conflict, and reticulate evolution among nuclear gene trees in *Oryza*. **a** Patterns of concordance and conflict between gene trees and the species tree, based on PhyParts analysis. The species tree was inferred with ASTRAL-III from 3,973 single-copy orthologous genes. For each branch, the top number indicates the count of concordant gene trees, and the bottom number indicates the count of conflicting trees. Pie charts show the proportion of concordant (blue), major conflicting (green), and other conflicting (red) topologies for major clades. **b** Phylogenetic network inferred from nuclear gene trees using SplitsTree, highlighting three major reticulate regions (colors correspond to clades in panel a). **c** The best-fit phylogenetic network with reticulation events inferred by PhyloNet under a MPL framework. Curved blue and red branches represent inferred hybridization events, with annotated inheritance probabilities (gamma). **d** Comparison of total log-likelihood scores for network models with 0 to 3 reticulation events.

A phylogenetic network reconstructed using SplitsTree visualized these pervasive conflicts, revealing clear reticulate structures between the FF and GG clades, between the BB and CC clades, and within the AA genome group (Fig. 2b), indicating a history shaped by non-tree-like processes.

Model-based network inference using PhyloNet identified two significant hybridization events as the best-fit model (Fig. 2c), which had a substantially higher log likelihood than alternatives (Fig. 2d, Additional file 1: Table S4). The model suggested: 1) deep introgression within the ancestral AA genome lineage (∼15.5% ancestry from a ghost lineage), and 2) introgression into the common ancestor of the FF and GG genomes from another ghost lineage (gamma = 0.013). These results indicate gene flow at multiple evolutionary stages has substantially shaped genome divergence.

### Quantifying the sources of discordance: introgression dominates over ILS

#### Evidence from allele frequencies

TreeMix analysis inferred an ancient gene flow event between *O. brachyantha* (FF) and *O. punctata* (BB) (Fig. 3a, Additional file 2: Fig. S3). Genome-wide *D*-statistics (ABBA-BABA tests) revealed pervasive introgression, with 181 of 220 tested quartets showing significant signals (Fig. 3b, Additional file 1: Table S5). Introgression proportions ranged from 0.07% to 46.22%, with notably high levels between closely related taxa like *O. sativa* and *O. nivara* (46.22%).

**Fig. 3.**
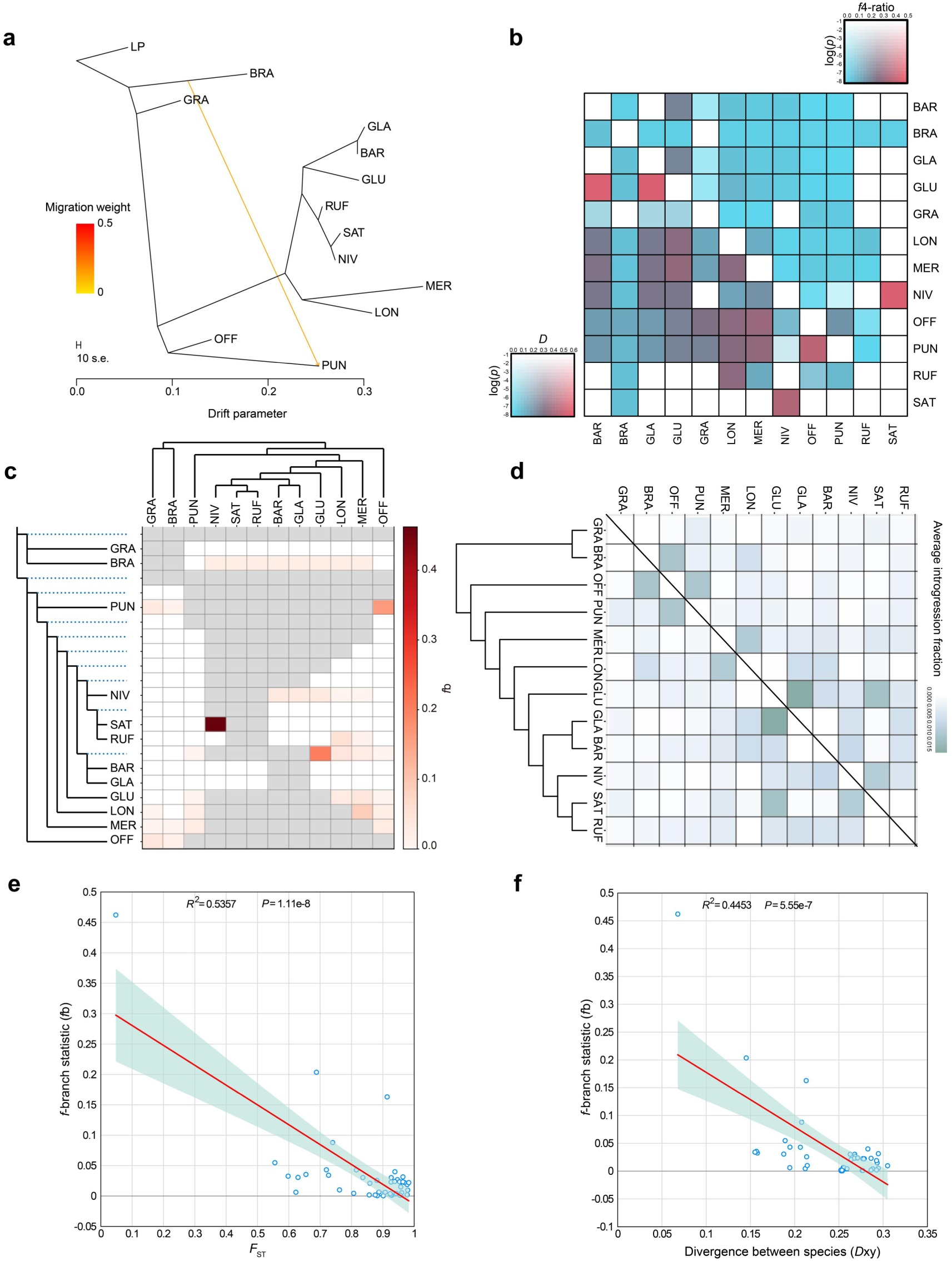
Quantifying introgression and its dominance over ILS in diploid *Oryza*. **a** Population splits and ancient gene flow inferred by TreeMix. Arrows indicate the direction and weight (color) of migration edges. **b** Heatmap of *D*-statistics (lower left) and *f*_4_-ratio values (upper right) calculated using Dsuite. **c** Results of the *f*b analysis, which quantifies branch-specific introgression. Each row in the matrix corresponds to a branch on the species tree (left), with color intensity reflecting the introgression proportion (*f*b). **d** Species-level total introgression proportion (totalIntroProp) inferred by QuIBL, a model-comparison method that distinguishes ILS from introgression based on gene tree branch lengths. **e** Negative correlation between the *f*b and *F*_ST_. **f** Negative correlation between *f*b and *D*xy. Species abbreviations: LP, *L. perrieri*; BRA, *O. brachyantha*; GRA, *O. granulata*; GLA, *O. glaberrima*; BAR, *O. barthii*; GLU, *O. glumaepatula*; RUF, *O. rufipogon*; SAT, *O. sativa*; NIV, *O. nivara*; MER, *O. meridionalis*; LON, *O. longistaminata*; OFF, *O. officinalis*; PUN, *O. punctata*.

#### Branch-specific signals

The *f*b statistic, which maps introgression signals onto species tree branches, revealed excess allele sharing along multiple internal branches, involving FF-BB/CC, GG-BB/CC, BB-CC, and CC-AA comparisons (Fig. 3c, Additional file 1: Table S6). As expected under gene flow, fb values were negatively correlated with genetic divergence (*F*_ST_) (*R*^2^ = 0.536, *P* = 1.11 x 10^-8^; Fig. 3e, Additional file 1: Table S7).

#### Model-based quantification (QuIBL)

To directly quantify the relative contributions, we used QuIBL. Of 360 discordant trios, 267 (74.17%) strongly supported the introgression model (Delta BIC < −10), while only 56 (15.56%) supported the ILS-only model. Finally, 30 trios (8.33%) had Delta BIC values between 0 and −10. As these values fall below the significance threshold, they are attributed to ILS or other stochastic processes rather than to gene flow. (Fig. 3d, Additional file 1: Table S8). The remaining 7 trios were excluded for not satisfying the topological assumptions. This indicates post-speciation introgression is the primary driver of gene tree discordance in diploid *Oryza*. Introgression was pronounced within the AA-genome group and between FF and CC genomes, whereas gene flow between AA and other genomes was more limited (Additional file 1: Table S9).

Together, these results demonstrate phylogenetic discordance is shaped predominantly by introgression, with ILS as a secondary component.

### Genomic localization and functional enrichment of introgressed regions

Having established introgression as the dominant source of conflict, we localized introgressed genomic regions for six species trios with strong *D*-statistic signals. We calculated *f*_dM_, a statistic sensitive to blocks of allele sharing that, under the null hypothesis, shows a symmetric distribution around zero (Additional file 2: Fig. S4). Windows with the highest *f*_dM_ values, cumulatively matching the genome-wide introgression proportion estimated by the *f*_4_-ratio, were defined as candidate introgressed regions (Additional file 1: Table S5, Table S10). Candidate introgressed regions consistently showed a significant negative correlation between interspecific *D*xy and the estimated proportion of introgression between the two recipient lineages (P2 and P3) (Fig. 3f, Additional file 1: Table S11), supporting their origin via gene flow rather than ILS. To further reduce potential confounding effects, we excluded windows with interspecific *D*xy values above the genome-wide average and filtered out regions with anomalous sequencing depth. After these filtering steps, the final introgressed tracts varied in size (approximately 3.3-4.2 Mb) and gene content (approximately 1.3%-50.3% of annotated genes) across trios (Additional file 1: Table S12). This filtering step was applied to minimize potential biases introduced by regions with elevated interspecific *D*xy, which may not be fully compatible with expectations under recent introgression.

GO enrichment analysis revealed that introgressed genes are non-randomly associated with specific functions (Fig. 4, Additional file 1: Table S13, Additional file 2: Fig. S5-S7). Enriched terms included secondary metabolism, stress response, developmental processes, and organelle functions, with specific patterns varying among trios (e.g., RUF_SAT_NIV: development/stress; RUF_GLA_GLU: secondary metabolism). These patterns suggest gene flow may have contributed to adaptive evolution and lineage diversification.

**Fig. 4.**
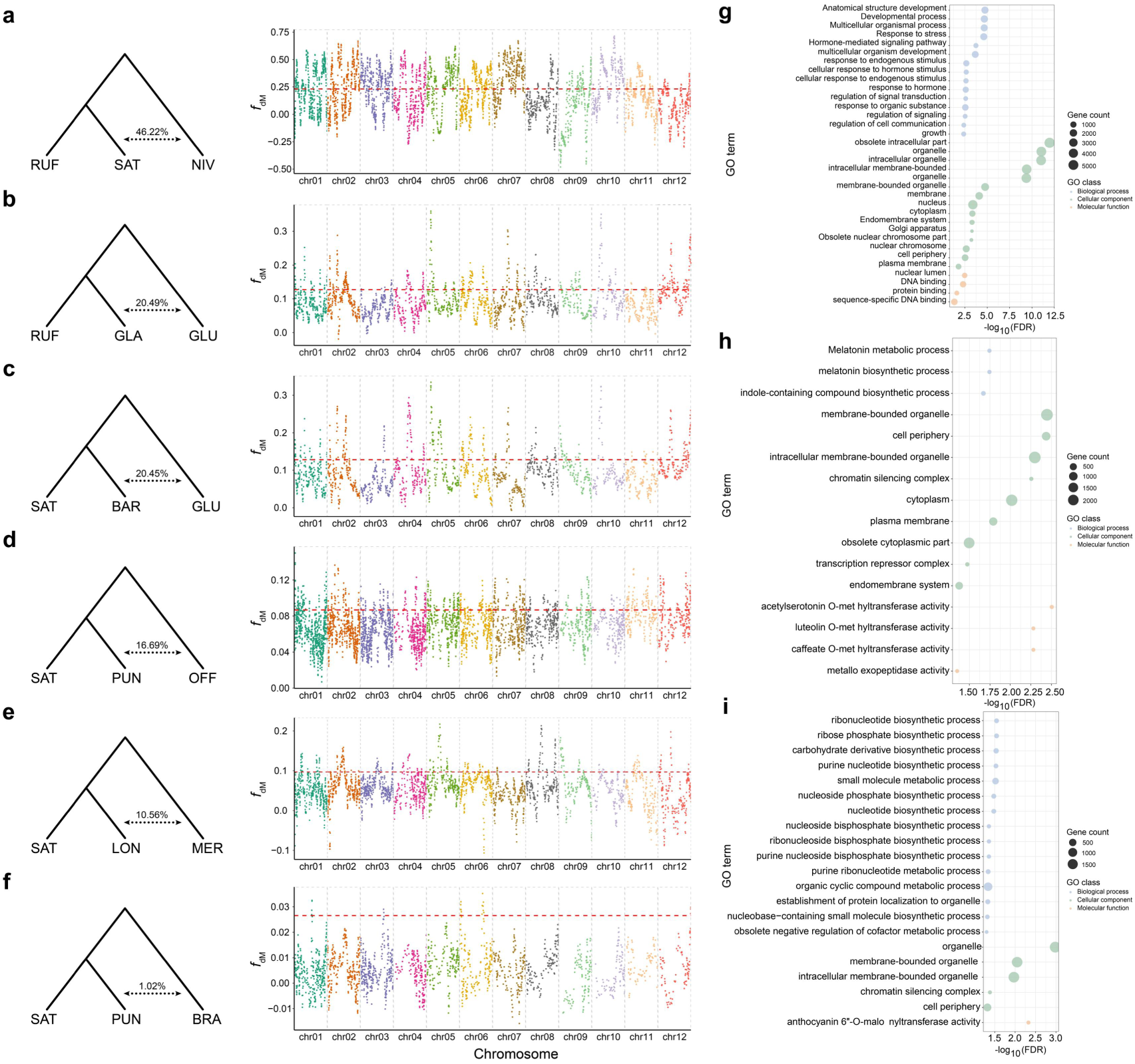
Genomic localization and functional enrichment of candidate introgressed regions. **a-f** For each of six species trios: the inferred trio topology with the genome-wide introgression proportion (*f*_4_-ratio), and a Manhattan plot of *f*_dM_ statistic values across the genome. The horizontal dashed line indicates the threshold for defining candidate introgressed windows. **g-i** GO enrichment analysis (FDR < 0.05) for genes within candidate introgressed regions for three representative trios: **g** RUF_SAT_NIV, **h** RUF_GLA_GLU, and **i** SAT_BAR_GLU. Bubble size represents the number of genes, and color indicates the functional category. Results for the remaining trios (SAT_PUN_OFF, SAT_LON_MER, SAT_PUN_BRA) are in Figures S9-S11.

### Divergence time estimation

Using the obtained robust phylogeny and calibrating with the *O. meridionalis* divergence time (∼2.32 Ma; TimeTree database), we estimated divergence times within diploid *Oryza* (Fig. 5). The estimated times for AA-genome species were consistent with a previous study [50]. Key findings include: the split between the FF/GG ancestor and the EE lineage at ∼13.48 Ma (Miocene), the divergence of GG and FF genomes at ∼12.21 Ma, and the very recent divergence of the two GG species (*O. granulata* and *O. meyeriana*) at ∼0.3 Ma (Pleistocene) (Fig. 5). The diversification of the AA genome lineage (∼2.3 Ma onward) coincides with a period of pronounced glacial-interglacial cycles, suggesting repeated climatic fluctuations may have promoted geographic fragmentation and reproductive isolation, contributing to its rapid diversification.

**Fig. 5.**
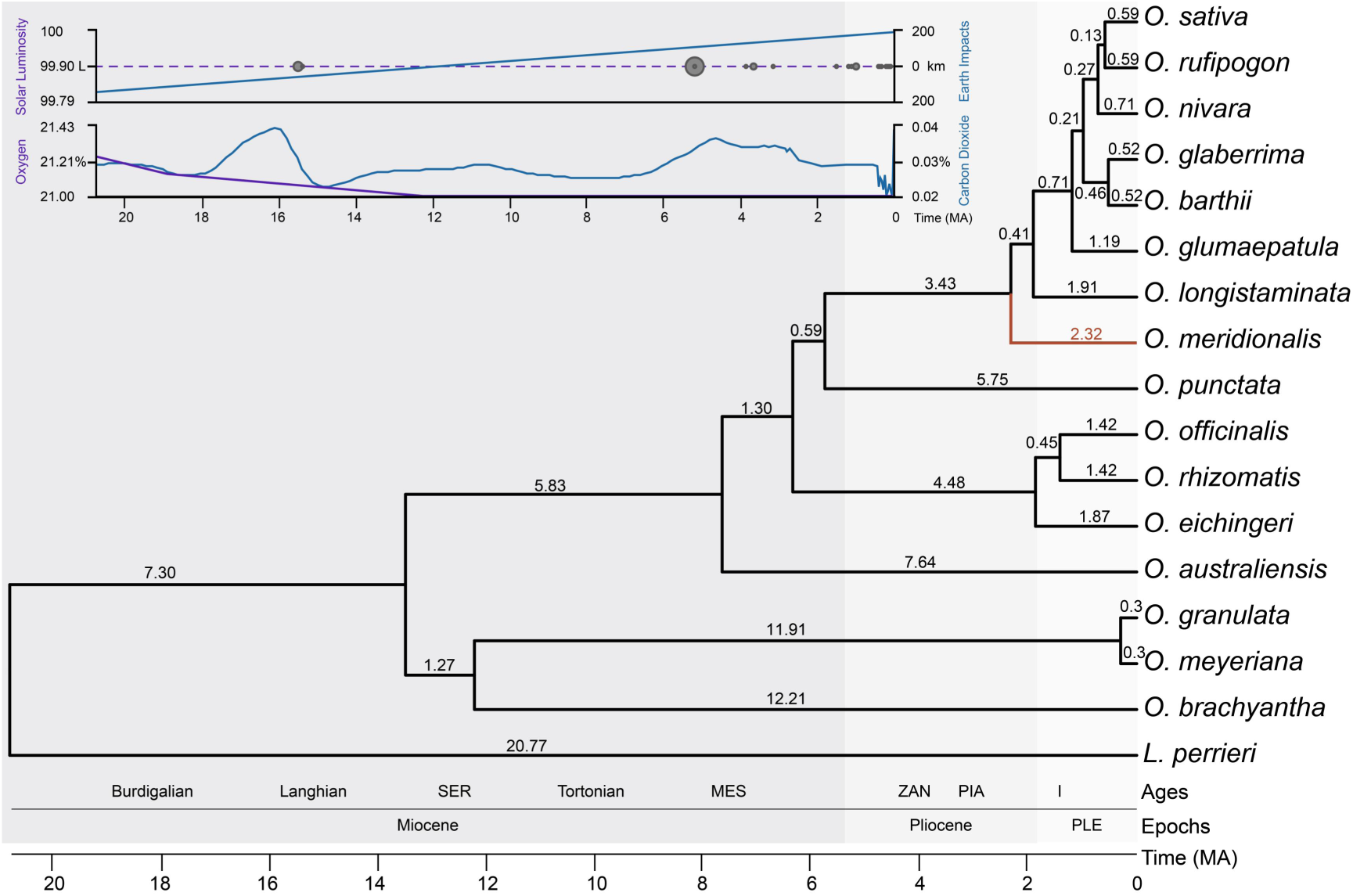
Divergence time estimation for diploid *Oryza* lineages in a paleoclimatic context. The chronogram was estimated using the transcriptome-based phylogeny, calibrated with the *O. meridionalis* divergence time (∼2.32 Ma). Node ages are shown with blue bars representing 95% confidence intervals. The background shows a schematic paleotemperature curve over the last 20 million years for reference.

## Discussion

### A robust phylogeny and pervasive discordance in *Oryza*

This study reconstructs a robust phylogenetic framework for diploid *Oryza* by integrating nuclear genome SNPs, single-copy orthologous genes from both genome and transcriptome datasets, and complete chloroplast genome sequences. Multiple analytical approaches including concatenated ML, coalescent-based species tree inference (ASTRAL), and analyses of gene tree topology frequencies consistently recovered a topology in which the FF and GG genome lineages form a monophyletic basal clade, sister to a clade containing the EE, CC, BB, and AA genomes (Fig. 1). This topology, strongly supported across independent datasets, differs from some earlier studies that proposed sequential divergence of GG then FF [23, 51, 59]. The high congruence across data types and the substantial proportion of individual gene trees (∼40%) supporting the FF+GG clade confirm this as a stable phylogenetic feature.

Despite this stable backbone, we detected extensive gene tree discordance. PhyParts analysis of 3,973 gene trees revealed a high frequency of alternative topologies at key nodes, particularly those associated with the early diversification of the CC, BB, and AA genome lineages. SplitsTree analyses further visualized these pervasive conflicts as network-like patterns among the FF/GG lineages, between the BB/CC lineages, and within the AA genome clade (Fig. 2b), suggesting underlying reticulate processes. Additionally, cytonuclear discordance within the AA lineage, where chloroplast data place *O. meridionalis*, not *O. longistaminata*, as the earliest-diverging lineage, provides classic evidence for historical chloroplast capture via hybridization. Thus, the evolutionary history of *Oryza* is characterized by a well-supported species tree overprinted by widespread genealogical heterogeneity.

### Introgression as the dominant driver of phylogenetic conflict

Discordance between gene trees and the species tree can arise from two major processes: introgressive hybridization and ILS [4]. Introgression introduces mosaic genomic segments via interspecific gene flow, while ILS arises from the stochastic sorting of ancestral polymorphisms during rapid successive speciation [68]. Disentangling their relative contributions requires an integrative framework. Our multi-pronged approach consistently identified historical gene flow as the predominant factor.

Signals of introgression were widespread. TreeMix inferred an ancient gene flow event between *O. brachyantha* (FF) and *O. punctata* (BB) (Fig. 3a). Genome-wide *D*-statistics revealed significant allele sharing in over 80% of tested quartets, with introgression proportions ranging up to 46.22% (Fig. 3b). The *f*b statistic mapped significant gene flow to specific internal branches across the phylogeny (Fig. 3c). These patterns are pronounced within the AA genome clade, aligning with documented historical sympatry and gene flow following the introduction of Asian rice to Africa [72–74], including evidence of introgression from *O. sativa* into *O. barthii* [75].

Critically, model comparison using QuIBL provided a direct, quantitative partition of the two processes [48]. This analysis demonstrated that approximately 74.17% of gene tree discordance is best explained by post-speciation introgression, with only about 15.56% attributable to ILS alone (Fig. 3d). This conclusion is reinforced by the non-random genomic distribution of introgression signals identified by *f*_dM_ scans; these signals form discrete blocks with reduced interspecific divergence, a pattern inconsistent with genome-wide ILS. Therefore, while ILS contributes, hybridization and introgression are the principal architects of phylogenetic incongruence in diploid *Oryza*.

### The value of an integrative phylogenomic framework

Beyond its specific conclusions about *Oryza*, this study demonstrates the power of an integrative, multi-method framework to dissect the sources of phylogenetic discordance. No single method is sufficient under all scenarios. We leveraged the complementary strengths of several approaches: 1) Species Tree Estimation (ASTRAL): Provided a robust topological hypothesis accounting for ILS [76], serving as an essential reference; 2) Network-Based Methods (SplitsTree, PhyloNet): Offered visual and model-based evidence for reticulation [77, 78], identifying specific hybridization events; 3) Allele-Frequency Statistics (*D*-statistic, *f*-branch): Gave intuitive, branch-specific tests for the direction and magnitude of introgression [46, 79]; 3) Model-Based Comparison (QuIBL): Enabled the key quantitative step of partitioning discordance between ILS and introgression models based on gene tree branch length distributions [48]; and 4) Genomic Scans (*f*_dM_): Allowed fine-scale localization of introgressed tracts [47], linking population genetic signals to specific genomic regions.

The strong concordance among results from these independent lines of evidence underscores the robustness of our conclusions. This framework is generalizable to other lineages facing challenges from rapid radiation and reticulation. Future advances, particularly with improved models for polyploid analyses and pangenome references, will further enhance such integrative studies.

## Conclusions

In summary, by applying an integrative phylogenomic framework to extensive data from diploid *Oryza*, we have achieved two main outcomes. First, we resolved a stable species phylogeny with FF and GG genomes forming a basal clade, diverging from other lineages during the Miocene (∼13.5 Ma). Second, and more broadly significant, we quantitatively demonstrated that historical introgression, not incomplete lineage sorting, is the predominant force generating genome-wide gene tree discordance in this genus. The introgressed genomic regions are enriched for genes involved in stress response, metabolism, and development, suggesting adaptive consequences of this widespread gene flow. Ultimately, this work exemplifies how combining complementary phylogenomic methods can successfully disentangle complex evolutionary histories, offering a methodological blueprint for studying rapid radiations and reticulate evolution.

## Methods

### Plant materials and data sources

Plant materials for 16 diploid *Oryza* species were grown under uniform greenhouse conditions. Total RNA was independently extracted from fresh, fully expanded leaves for transcriptome sequencing with three replicates per species, and low-coverage whole-genome resequencing (∼10 x depth) for two accessions per species, which were generated by our laboratory and have been previously published [80]. To augment our data, six additional resequencing datasets and one transcriptome dataset were downloaded from public databases. Detailed information on all resequencing samples is provided in Additional file 1: Tables S1. In addition to the sequencing data generated here, we utilized publicly available genome assemblies from 12 diploid *Oryza* species.

Protein sequences for ten *Oryza* species and the outgroup *L. perrieri* were downloaded from the EnsemblPlants database (https://plants.ensembl.org/info/data/ftp/index.html). These included *O. sativa* ssp. *japonica* cv. ‘Nipponbare’, *O. nivara*, *O. glaberrima*, *O. barthii*, *O. glumaepatula*, *O. meridionalis*, *O. rufipogon*, *O. longistaminata*, *O. punctata*, and *O. brachyantha*. Protein sequences for *O. granulata* [81, 82] and *O. officinalis* [83] were obtained from their respective genome project websites.

### Whole-genome resequencing and single-nucleotide polymorphism (SNP) calling

Genomic DNA was extracted using a modified CTAB method [84]. Following quality assessment, DNA was fragmented by ultrasonication, and sequencing libraries were constructed through standard steps of end repair, A-tailing, adapter ligation, and size selection. Libraries were sequenced on an Illumina platform.

Bioinformatic processing of resequencing data was performed as follows. Raw reads were quality-filtered using Fastp v.0.21.0 with parameters -q 30 -u 40 -l 50 [85]. Quality-controlled reads were aligned to the *O. sativa* cv. ‘Nipponbare’ reference genome (IRGSP-1.0) using BWA. SAM files were converted, sorted, and processed using SAMtools v.1.1. PCR duplicates were identified and marked using MarkDuplicates in GATK v.4.2.0.0 and excluded from downstream analyses. Variants calling was performed per sample using GATK HaplotypeCaller with the --emit-ref-confidence GVCF mode, and individual GVCF files were merged and jointly genotyped. SNPs were extracted and subjected to preliminary hard filtering using VariantFiltration with the following criteria: QD < 2.0, MQ < 40.0, FS > 60.0, SOR > 3.0, MappingQualityRankSum < −12.5, or ReadPosRankSum < −8.0.

To obtain a high-confidence SNP set from conserved regions, we used OrthoFinder v.2.5.2 [86] to identify 4,266 single-copy genes from 12 available *Oryza* genomes.

SNPs within these gene regions were extracted using BEDTools v.2.29.2. Further stringent filtering was applied using BCFtools v.1.10.2 and VCFtools v.0.1.16, removing SNPs that met any of the following conditions: (1) read depth > 50 or < 2; (2) missing rate > 10%; (3) non-biallelic; (4) minor allele frequency (MAF) < 0.05; (5) more than three SNPs within a 10 bp window; or (6) located within 5 bp of an indel.

### Identification of single-copy orthologous genes

We identified single-copy orthologs from two complementary data sources to ensure robustness. First, a total of 3,973 single-copy orthologous genes were identified from 13 diploid genomes using OrthoFinder v.2.5.2 with parameter -S blast. Second, 846 single-copy orthologous genes were identified from 17 assembled transcriptomes (16 diploid *Oryza* species and *L. perrieri*; Additional file1: Table S3) using the same OrthoFinder pipeline. For phylogenetic analysis, each set of orthologous genes was aligned with MUSCLE v.3.8.31, and conserved regions were selected using Gblocks v.0.91b with parameters: -b4=10 -b5=n -t=p. Complete chloroplast genome sequences were also assembled for phylogenetic analysis.

### Phylogenetic analysis

Phylogenetic relationships were reconstructed from four distinct datasets using both concatenation-based and coalescent-based approaches to ensure topological robustness.

#### Phylogeny based on whole-genome resequencing SNPs

VCF files containing 187,135 SNPs at fourfold degenerate transversion (4DTv) sites and 209,888 SNPs in single-copy gene regions were converted to PHYLIP format using vcf2phylip.py (https://github.com/edgardomortiz/vcf2phylip). Maximum likelihood (ML) phylogenies were inferred with RAxML v.8.2.12 under the GTRGAMMA model [87] using 1000 bootstrap replicates (-x 12345 -p 12345 -# 1000). Trees were visualized with iTOL [88].

#### Phylogeny based on transcriptome single-copy orthologs

ML trees were constructed from the 846-gene concatenated alignment using RAxML v.8.2.12 with the PROTGAMMAIJTTF model and 1000 bootstrap replicates.

#### Phylogeny based on chloroplast genomes

To assess potential cytonuclear discordance indicative of historical introgression [35, 89–91], ML phylogenies were constructed from complete chloroplast genome alignments using RAxML with the same parameters as for nuclear SNPs.

#### Coalescent-based species tree and conflict assessment

From the 3,973 genomic single-copy orthologs, we inferred a species tree under the multispecies coalescent model using ASTRAL-III [76] on individual gene trees generated by RAxML. To visualize the extent of gene tree variation, we employed DensiTree [81] to visualize the topological variation among gene trees. Furthermore, to quantify gene tree concordance and conflict, we used PhyParts [92] to map all 3,973 gene trees onto the ASTRAL species tree. Gene trees were rooted consistently using the ETE Toolkit v.3.0.0S [93].

### Introgression and ILS analyses

To disentangle the contributions of introgression and ILS to gene tree discordance, we applied a suite of complementary methods designed to address specific questions within our integrative framework. These included phylogenetic network inference, allele-frequency-based analyses, and branch-length-based statistical tests: specifically, SplitsTree, PhyloNet, TreeMix, Patterson’s *D* statistic (ABBA-BABA test), the *f*-branch (*f*b) statistic, and QuIBL.

#### Phylogenetic Network Inference

A genome-wide set of 3,973 nuclear single-copy orthologous gene trees was used to infer a consensus network in SplitsTree, providing a visual representation of topological conflict. For model-based inference of specific reticulation events, we used the maximum pseudolikelihood (MPL) framework in PhyloNet v.3.8.2 [94, 95]. Networks with 0 to 3 reticulation events were inferred using the InferNetwork_MPL function. Ten independent searches were performed per reticulation level to avoid local optima. The optimal network, selected based on total log pseudo-likelihood, was visualized with IcyTree [96], and inheritance probabilities (gamma) for hybridization edges were estimated.

#### TreeMix Analysis

To infer the direction and magnitude of historical population-level gene flow, we used TreeMix v.1.13 [97] with OrientAGraph [98], specifying *L. perrieri* as the outgroup. Input files were prepared from 47,763 high-quality, independent SNPs obtained after stringent filtering (--max-missing 1 and --indep-pairwise 50 10 0.2). Models with 0 to 10 migration edges were each run ten times.

The optimal number of migration events was determined using the Evanno method implemented in the OptM package.

#### *D*-statistic and *f*-branch Analysis

To test for asymmetrical gene flow in specific taxon quartets, we calculated Patterson’s *D*-statistics using Dsuite [79] on 209,888 high-quality SNPs for all possible quartets among 12 taxa (220 combinations). This test, grounded in coalescent theory, evaluates allele frequency patterns (ABBA and BABA) to detect deviations from the null expectation of ILS alone [46, 99, 100]. A significant deviation from zero (|*Z*| > 3) indicates introgression. To localize introgression signals to specific branches of the species tree, we calculated the fb statistic, a phylogeny-based normalized metric (0 <= *f*b <= 1) which integrates *f*_4_-ratio estimates with the tree topology to quantify excess allele sharing between branches [79, 101].

#### QuIBL Analysis

As a core method to quantitatively distinguish and partition the effects of ILS versus introgression, we used QuIBL [48]. QuIBL applies a maximum likelihood framework to compare the distribution of internal branch lengths in gene trees under two competing models: a null model (H_0_) assuming discordance arises exclusively from ILS, and an alternative model (H_1_) incorporating both ILS and post-speciation introgression. Model fit was evaluated using the Bayesian Information Criterion (BIC), with Delta BIC < −10 considered strong support for the introgression model [102]. Analyses were conducted on all possible three-taxon combinations (trios) from the 12 species using the 3,973 gene trees. QuIBL parameters were: number of mixture distributions = 2, likelihood convergence threshold = 0.01, maximum steps = 50, gradient ascent scalar = 0.5. Note that QuIBL primarily detects post-speciation introgression [13].

### Identification of introgressed genomic regions

For species pairs with significant *D*-statistics, we performed genome-wide scans to localize introgressed regions. The fd and *f*_dM_ statistics [47, 103], which measure allele frequency patterns to identify genomic blocks with excess shared ancestry, were calculated in 100-kb sliding windows (10 kb steps). For each species triplet, candidate introgressed windows were defined as those with the highest *f*_dM_ values, cumulatively covering the genome-wide introgression proportion estimated by the *f*_4_-ratio [104].

To validate these candidates and distinguish true introgression from ILS, we calculated pairwise genetic divergence (*D*xy) for each candidate region using 100-bp windows and compared it to the chromosome-wide background *D*xy distribution (estimated in 50-kb windows using Pixy v.2.0.0.beta8) via the Mann-Whitney U test in R. Regions with *D*xy exceeding the chromosomal mean were excluded.

To control for mapping biases, sites with extreme sequencing depth (<5 or >40) were filtered using VCFtools. Finally, to explore the potential functional impact of introgression, genes within the validated introgressed regions were subjected to GO enrichment analysis using eggNOG-mapper v.2 [105] Significant terms were identified after Benjamini-Hochberg correction (FDR < 0.05).

### Divergence time estimation

Divergence times were estimated based on the robust phylogenetic framework obtained from the transcriptome-based single-copy orthologs, as these genes are less likely to be affected by introgression. The analysis was conducted using r8s v.1.80 under a relaxed molecular clock model. The divergence time of *O. meridionalis* was fixed at ∼2.32 Ma as a calibration point, based on the TimeTree database (which synthesizes 11 studies) and consistent with a recent genome-wide estimate of ∼2.41 Ma [50]. Branch lengths were optimized by maximum likelihood with default parameters.

### Overview of the integrative phylogenomic framework

To systematically resolve the sources of phylogenetic discordance in *Oryza*, we implemented an integrative analysis pipeline. This workflow sequentially addressed: (1) reconstruction of a robust species tree from multiple data types using concatenation and coalescent methods; (2) genome-wide assessment of gene tree conflict and visualization of reticulate signals; (3) discrimination and quantification of the relative contributions of ILS versus introgression to the observed conflict using complementary model-based and statistic-based approaches; (4) genomic localization and functional annotation of introgressed regions; and (5) estimation of divergence times within the resolved phylogenetic framework. A schematic overview of this analytical workflow is provided in Additional file 2: Figure S8.

## Supporting information

Supplemental Figure 1, and will be used for the link to the file on the preprint site.

Supplemental Figure 2, and will be used for the link to the file on the preprint site.

Supplemental Figure 3, and will be used for the link to the file on the preprint site.

Supplemental Figure 4, and will be used for the link to the file on the preprint site.

Supplemental Figure 5, and will be used for the link to the file on the preprint site.

Supplemental Figure 6, and will be used for the link to the file on the preprint site.

Supplemental Figure 7, and will be used for the link to the file on the preprint site.

Supplemental Figure 8, and will be used for the link to the file on the preprint site.

Supplemental Table 1, and will be used for the link to the file on the preprint site.

Supplemental Table 2, and will be used for the link to the file on the preprint site.

Supplemental Table 3, and will be used for the link to the file on the preprint site.

Supplemental Table 4, and will be used for the link to the file on the preprint site.

Supplemental Table 5, and will be used for the link to the file on the preprint site.

Supplemental Table 6, and will be used for the link to the file on the preprint site.

Supplemental Table 7, and will be used for the link to the file on the preprint site.

Supplemental Table 8, and will be used for the link to the file on the preprint site.

Supplemental Table 9, and will be used for the link to the file on the preprint site.

Supplemental Table 10, and will be used for the link to the file on the preprint site.

Supplemental Table 11, and will be used for the link to the file on the preprint site.

Supplemental Table 12, and will be used for the link to the file on the preprint site.

Supplemental Table 13, and will be used for the link to the file on the preprint site.

## Supplementary Information

**Additional file 1: Table S1.** Plant materials for re-sequencing and data sources. **Table S2.** Chromosome distribution of SNPs located in single copy orthologous gene region and 4DTv locus. **Table S3.** Summary of the assembled transcriptome of *Oryza* species. **Table S4.** Total Log-Likelihood Values from PhyloNET Analysis for the 12 Diploid *Oryza* Species. **Table S5.** Results of *D* and *f*4-ratio statistics from Dsuite analysis. **Table S6.** Number of shared alleles between *Oryza* populations based on *f*-branch analysis using Dsuite. **Table S7.** Introgression proportion (*f*b) vs. *F*_ST_ for species pairs. **Table S8.** Results of QuIBL analysis. **Table S9.** Average total introgression proportion per species pair in the QuIBL analysis. **Table S10.** Values of *f*d and *f*_dM_ based on the sliding window analysis of the six trios. **Table S11.** Introgression proportion (*f*b) vs. genetic differentiation (*D*xy) for species pairs. **Table S12.** Properties of introgressed regions in 6 trios. **Table S13.** Summary of significant GO terms across six trios (FDR < 0.05).

**Additional file 2: Figure S1.** Statistics and analysis of SNPs mapped to reference genome. **Figure S2.** Evaluation results of the assembled transcriptome by using BUSCO software. **Figure S3.** The optimal number of migration events (*m*) as predicted by OptM. **Figure S4.** Distribution of *f*_dM_ statistic values across all windows in the genome among the six trios (corresponding to Table S10). **Figure S5.** SAT_PUN_OFF trio GO enrichment of genes within candidate introgressed windows. **Figure S6.** SAT_LON_MER trio GO enrichment of genes within candidate introgressed windows. **Figure S7.** SAT_PUN_BRA trio GO enrichment of genes within candidate introgressed windows. **Figure S8.** An integrative phylogenomic framework for dissecting sources of genome-wide phylogenetic discordance.

## Acknowledgments

This work was supported by a start-up grant from Hainan University (to L.-Z.G.).

## Author Contributions

L.-Z.G. designed the study. H.-F.L., S.-F.D. and T.-L.F. analyzed the data. S.-F.D. performed the experiments. H.-F.L. drafted the manuscript. L.-Z.G. revised the manuscript.

## Funding

This work was supported by a startup grant of Hainan University to Li-zhi Gao.

## Declarations

### Ethics approval and consent to participate

Not applicable.

### Consent for publication

Not applicable.

### Availability of data and materials

The raw sequence data from this study are publicly available in the Genome Sequence Archive (GSA) at the China National Center for Bioinformation (CNCB)/National Genomics Data Center (NGDC). The transcriptome data are under BioProject PRJCA056805 (Additional file 1: Table S3). Accession details for all publicly referenced genome assemblies and transcriptomes of diploid *Oryza* species are provided in the Methods section.

### Competing interests

The authors declare that they have no competing interests.

## Notes

### Competing Interest Statement

The authors have declared no competing interest.

