## Supplementary figures and images for "An Integrative Phylogenomic Framework Quantifies the Dominant Role of Introgression in Phylogenetic Discordance among Diploid *Oryza* Species"

### Supplemental Figure 1, and will be used for the link to the file on the preprint site.

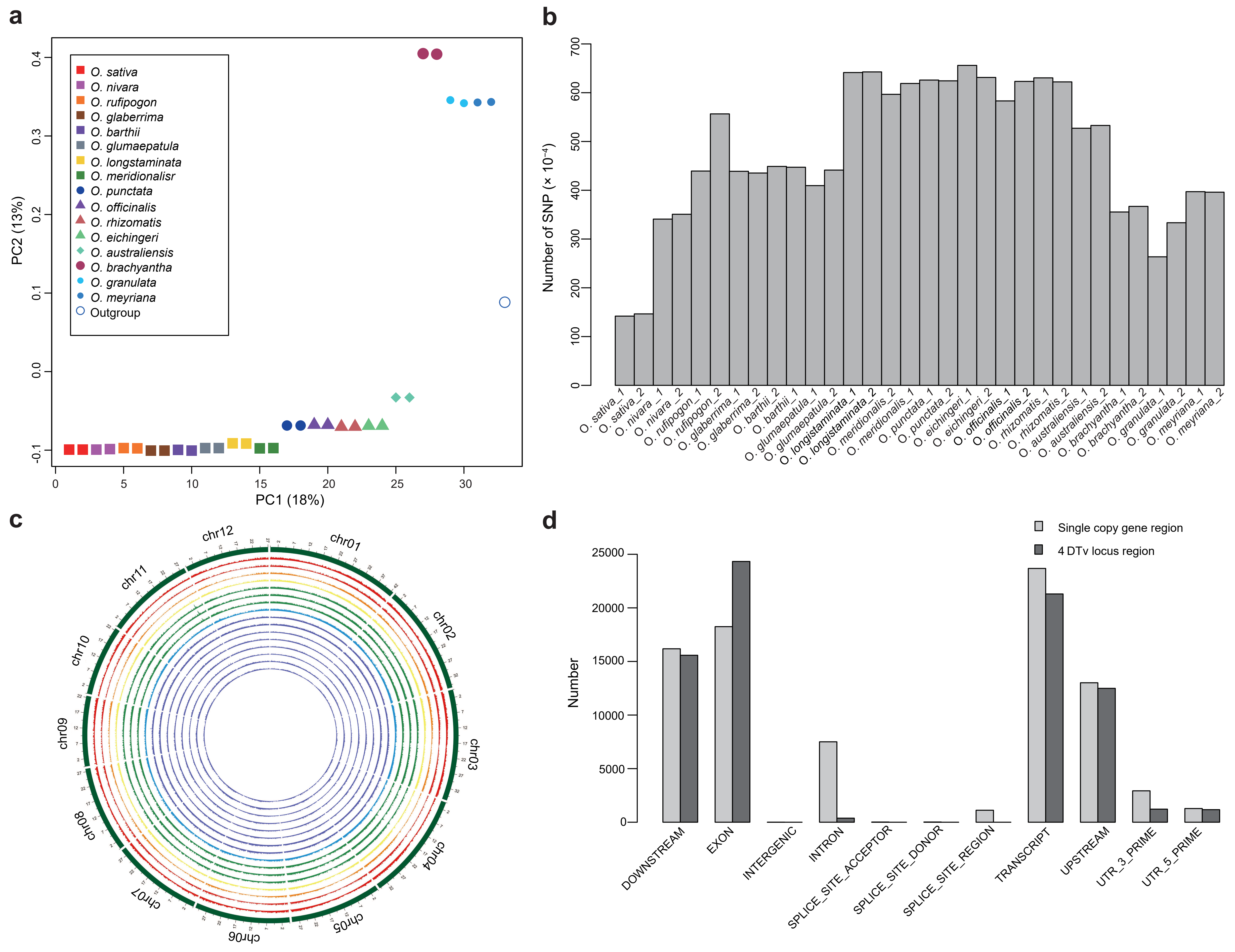

### Supplemental Figure 2, and will be used for the link to the file on the preprint site.

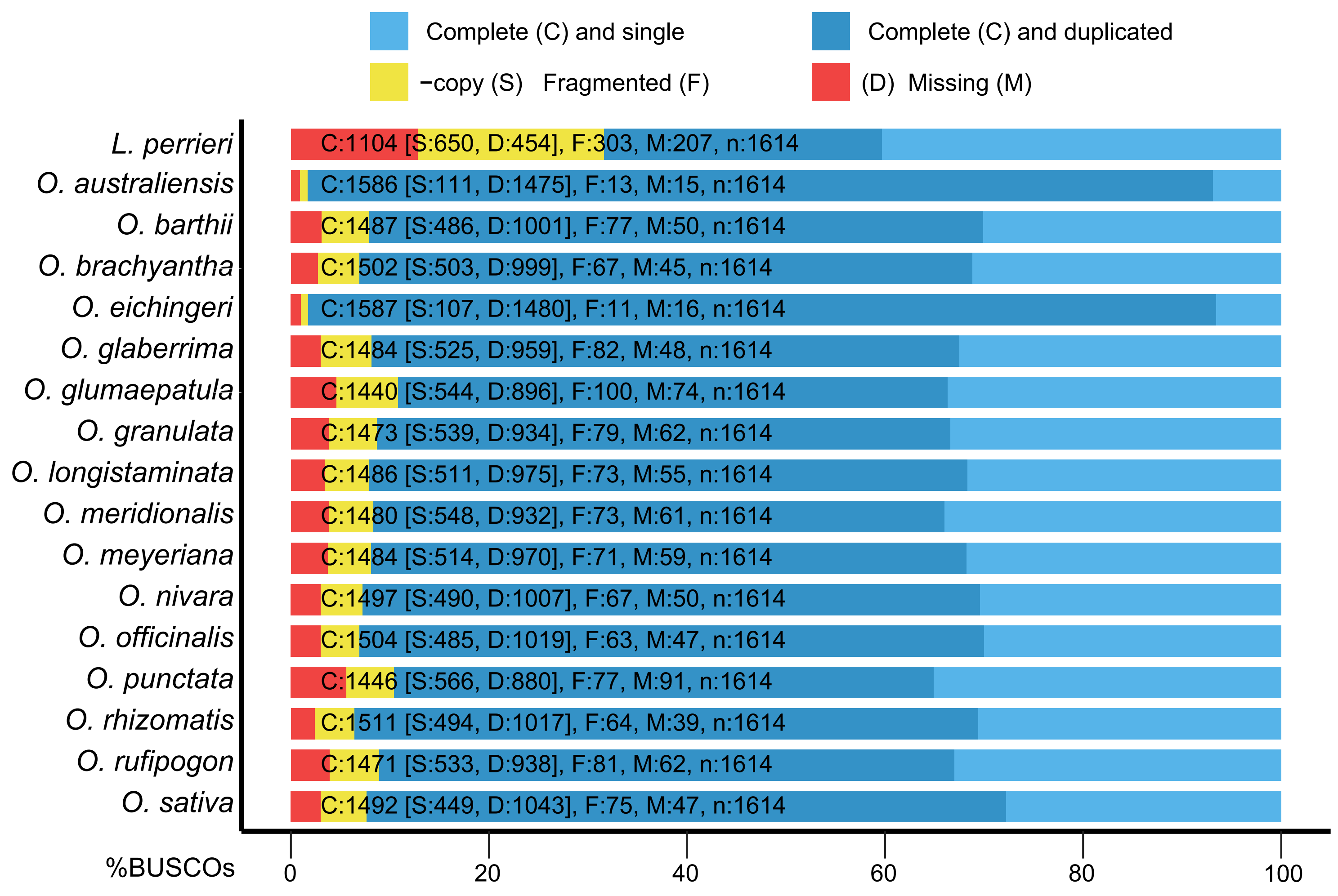

### Supplemental Figure 3, and will be used for the link to the file on the preprint site.

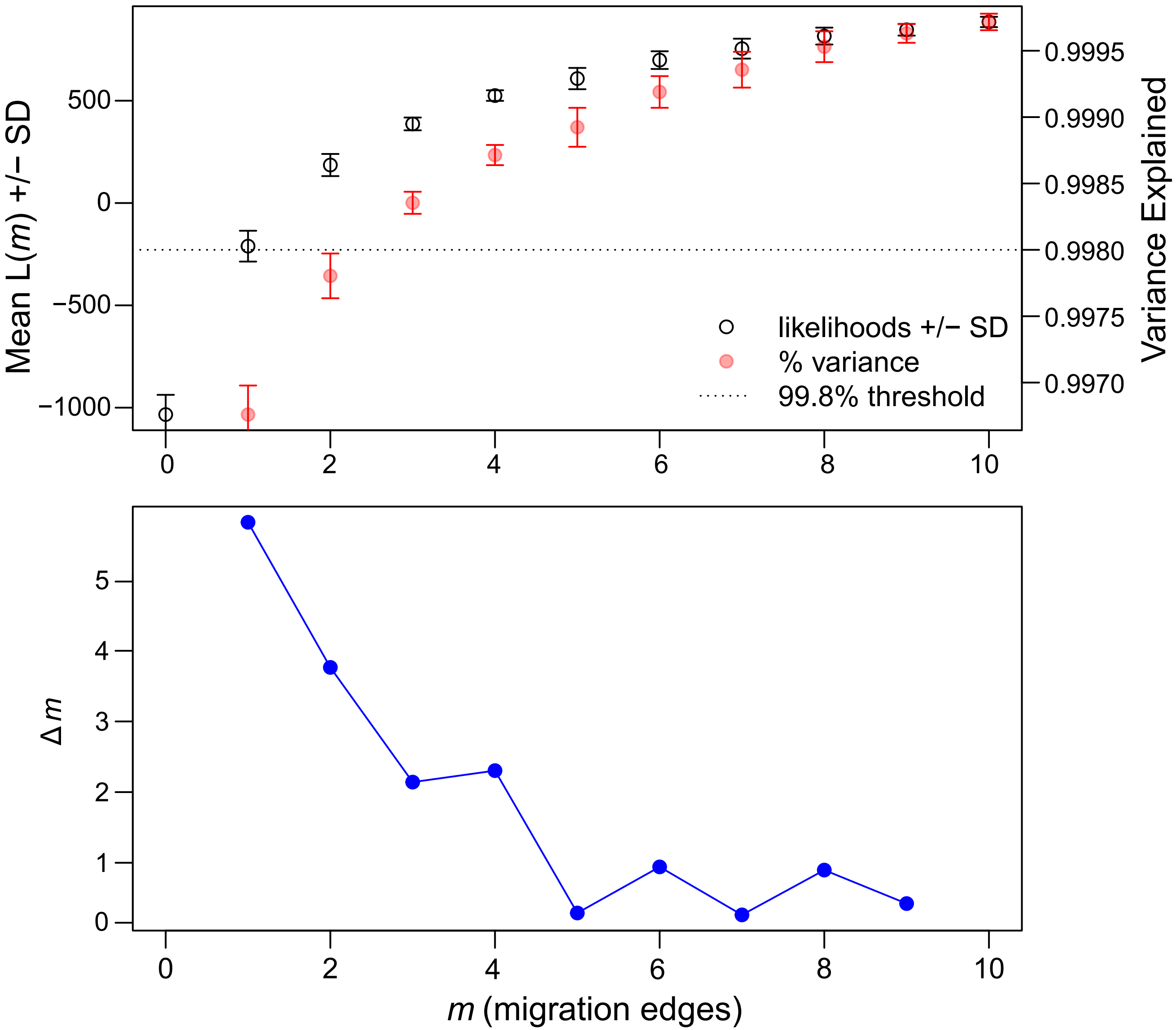

### Supplemental Figure 4, and will be used for the link to the file on the preprint site.

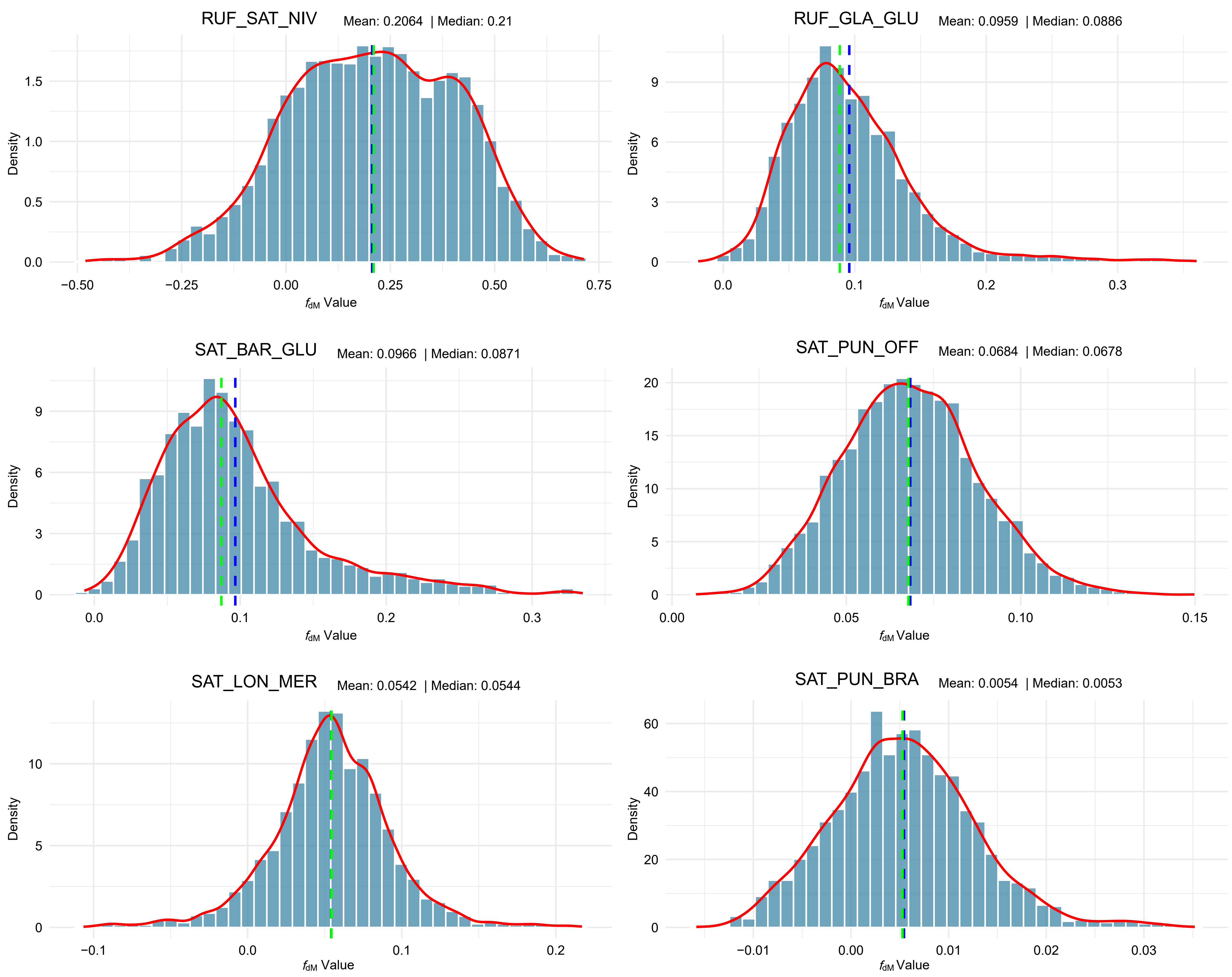

### Supplemental Figure 5, and will be used for the link to the file on the preprint site.

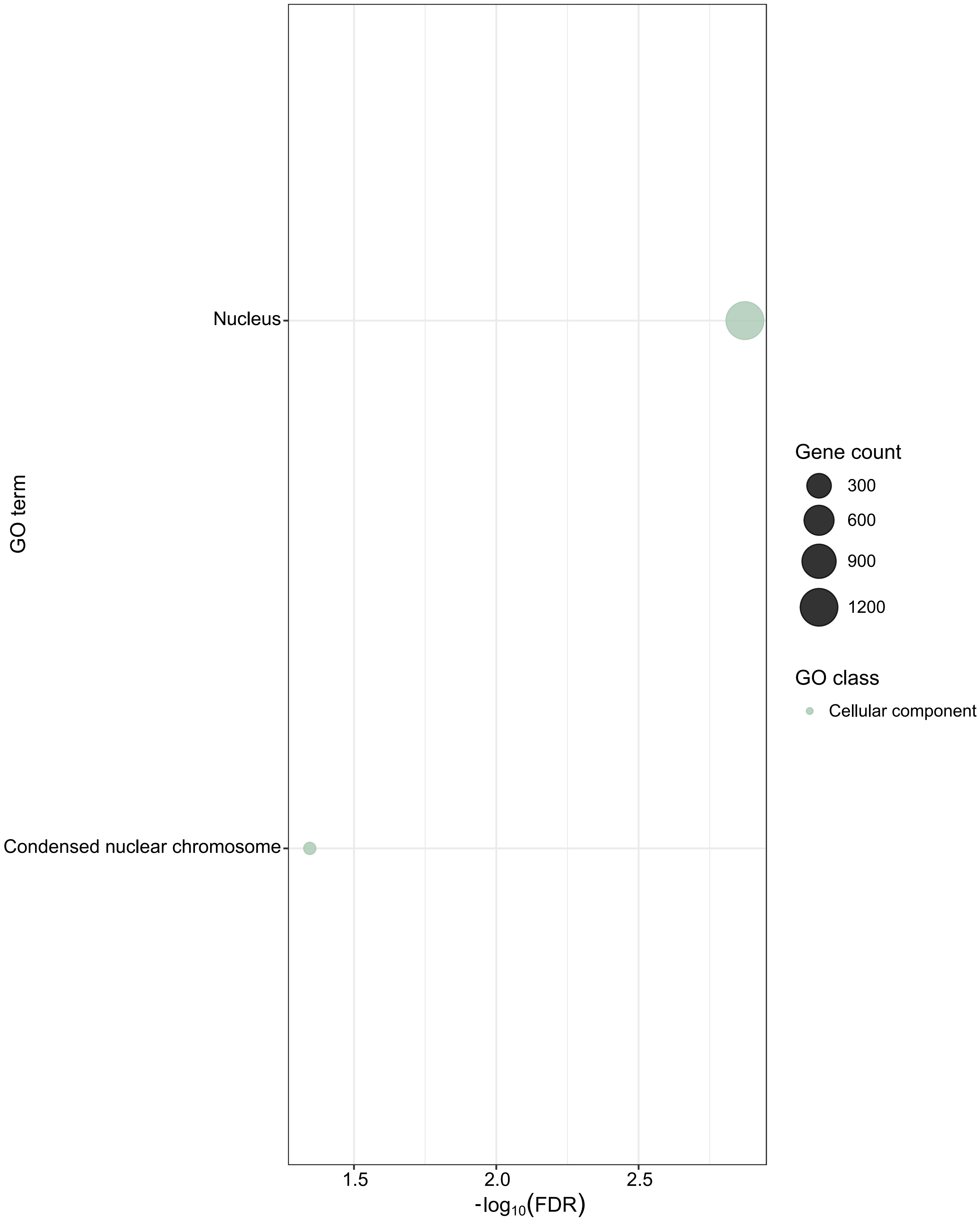

### Supplemental Figure 6, and will be used for the link to the file on the preprint site.

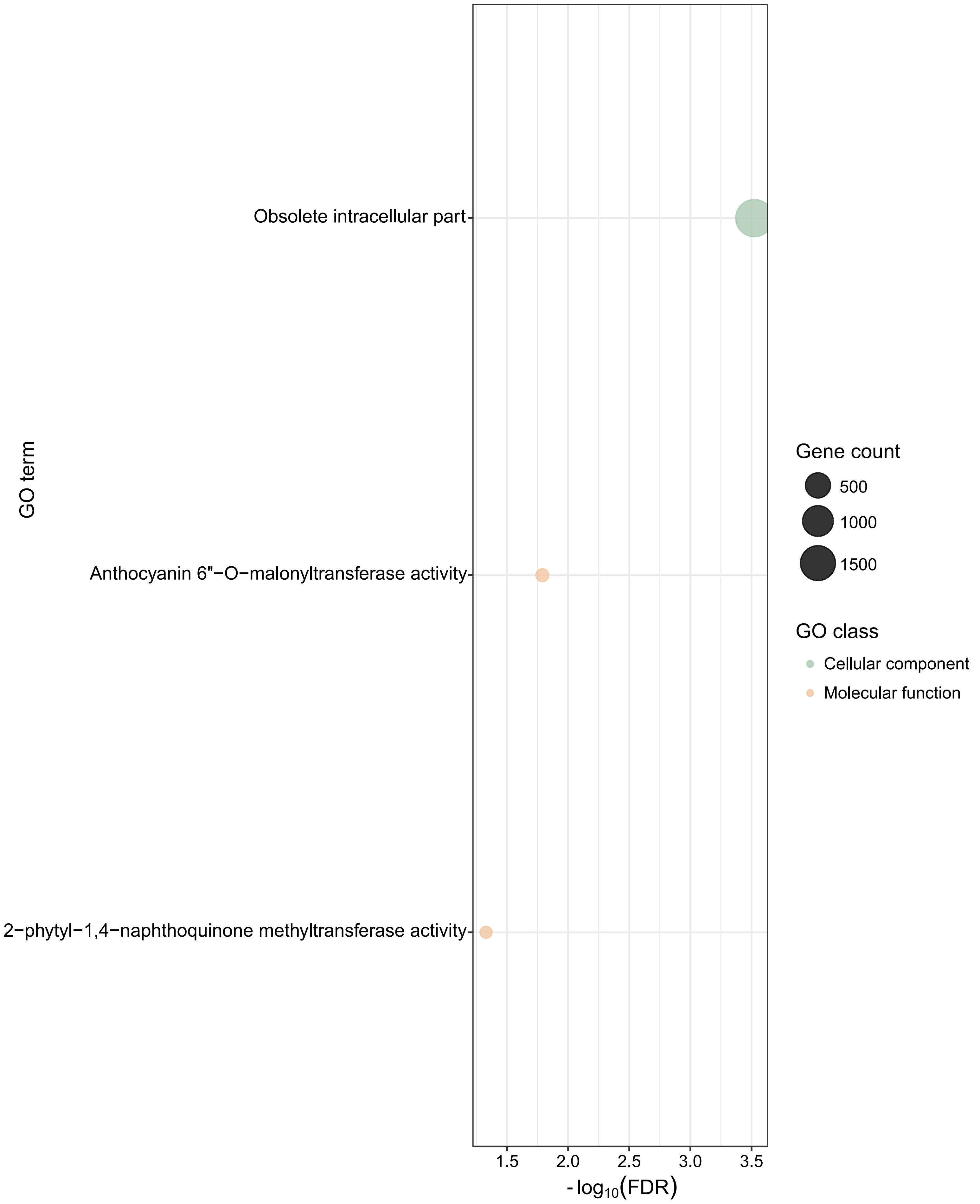

### Supplemental Figure 7, and will be used for the link to the file on the preprint site.

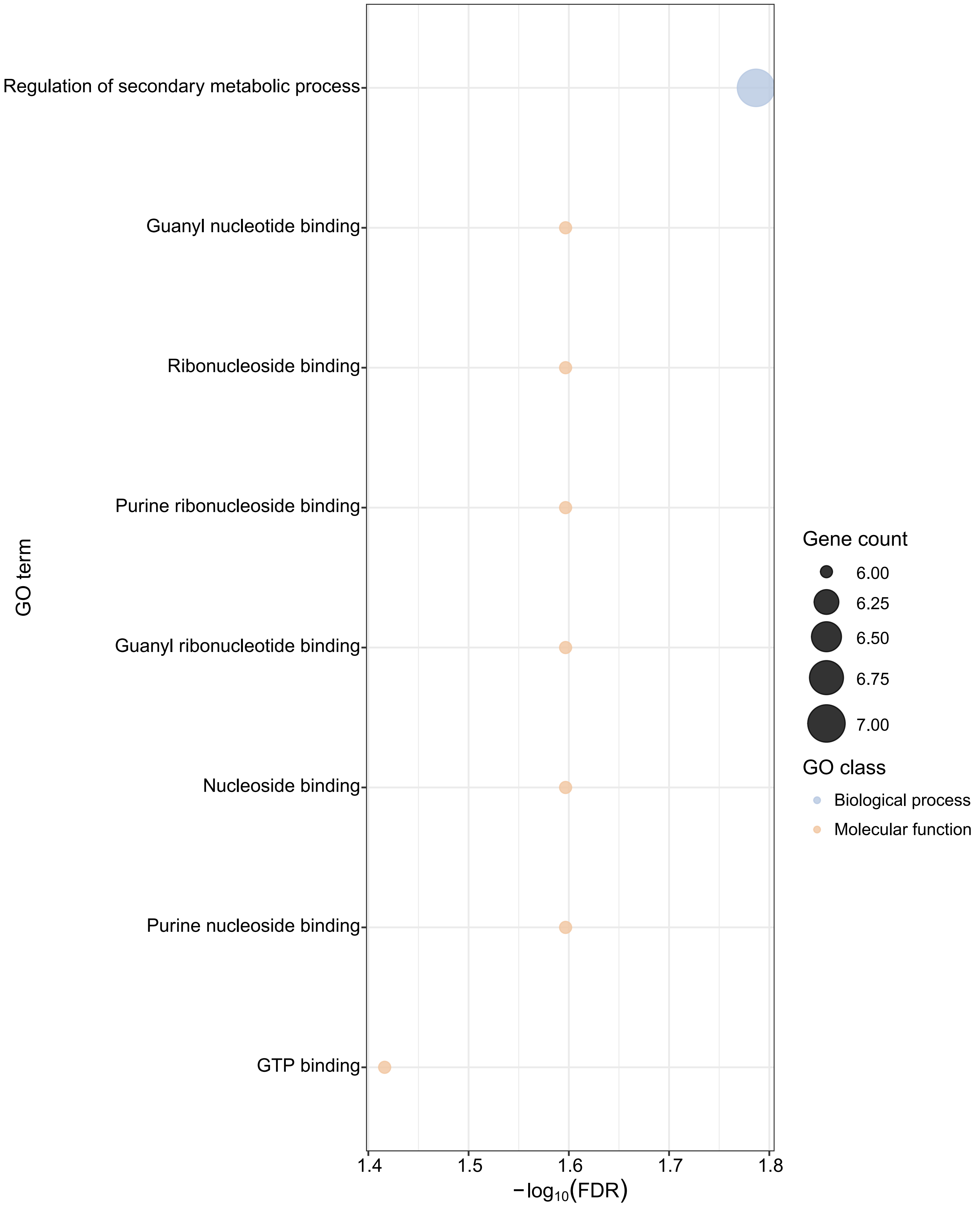

### Supplemental Figure 8, and will be used for the link to the file on the preprint site.

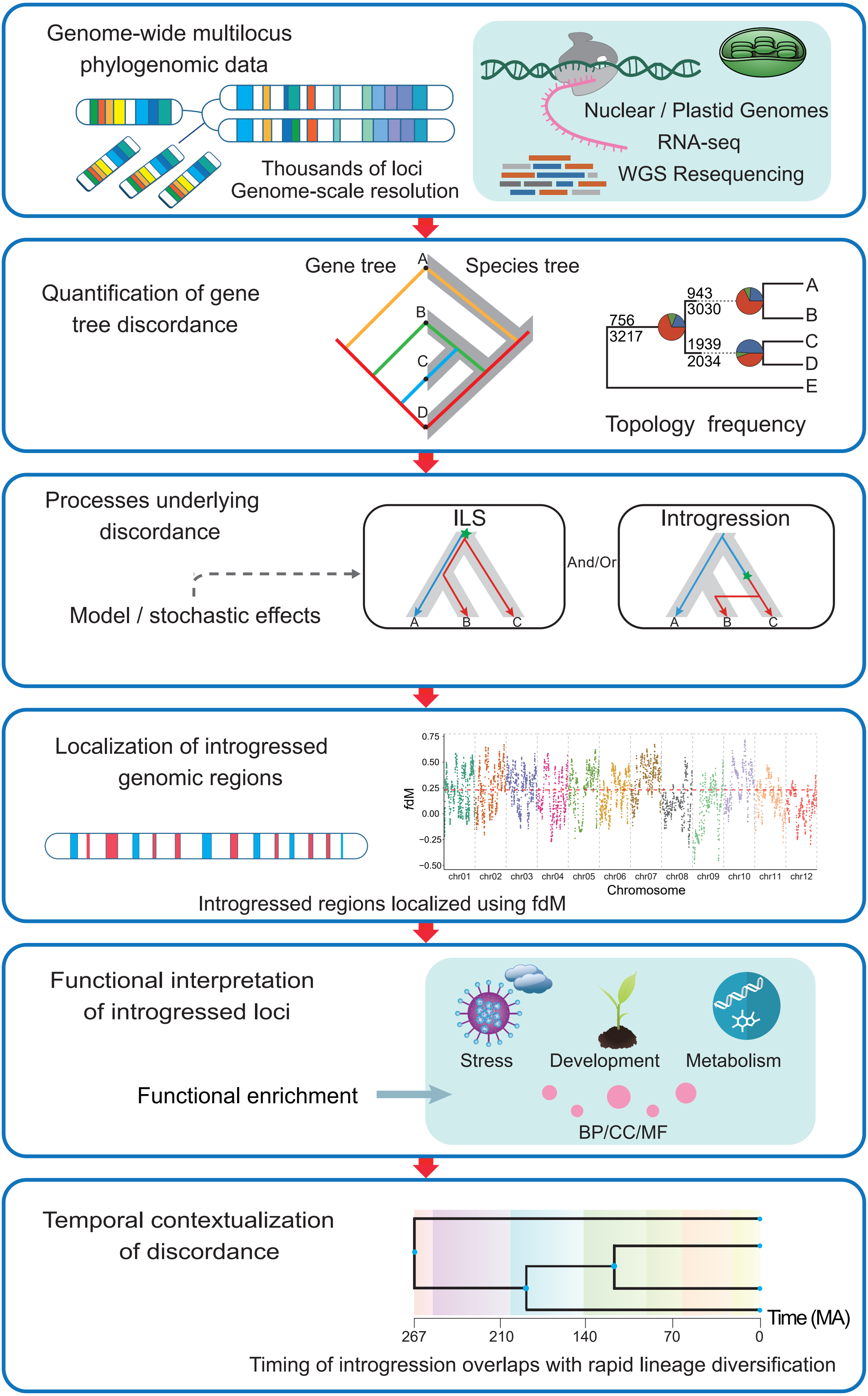
